# Spatial sorting as the spatial analogue of natural selection

**DOI:** 10.1101/210088

**Authors:** Ben L. Phillips, T. Alex Perkins

## Abstract

In most systems, dispersal occurs despite clear fitness costs to dispersing individuals. Theory posits that spatial heterogeneity in habitat quality pushes dispersal rates to evolve towards zero, while temporal heterogeneity in habitat quality favours non-zero dispersal rates. One aspect of dispersal evolution that has received a great deal of recent attention is a process known as spatial sorting, which has been referred to as a “shy younger sibling” of natural selection. More precisely, spatial sorting is the process whereby variation in dispersal ability is sorted along density clines and will, in nature, often be a transient phenomenon. Despite this transience, spatial sorting is likely a general mechanism behind non-zero dispersal in spatiotemporally varying environments. While generally transient, spatial sorting is persistent on invasion fronts, where its effect cannot be ignored, causing rapid evolution of traits related to dispersal. Spatial sorting is described in several elegant models, yet these models require a high level of mathematical sophistication and are not accessible to most evolutionary biologists or their students. Here, we frame spatial sorting in terms of the classic haploid and diploid models of natural selection. We show that, on an invasion front, spatial sorting can be conceptualized precisely as selection operating through space rather than (as with natural selection) time, and that genotypes can be viewed as having both spatial and temporal aspects of fitness. The resultant model is strikingly similar to classic models of natural selection. This similarity renders the model easy to understand (and to teach), but also suggests that many established theoretical results around natural selection could apply equally to spatial sorting.

## Introduction

Natural selection, the primary driver of evolutionary change, operates to maximise fitness. Because natural selection is such a powerful force, it is noteworthy when processes are unearthed that operate to displace a population from its fitness optimum. We are well aware of stochastic processes that undermine natural selection (mutation, drift, and its spatial analogue, the founder effect; Slatkin and Excoffier (2012)), but deterministic processes that act to reduce fitness are unusual. One such process is gene flow, in which maladapted alleles are continually introduced to a population (Lenormand 2002) by dispersing organisms (Ronce 2007). Dispersal is an interesting trait beyond simply causing gene flow, however. Models in which dispersal itself evolves often result in populations that are displaced from their fitness optimum, meaning that fitness is sacrificed to maintain dispersal.

The author of one of the earliest models of dispersal evolution was so struck by the fact that dispersal might evolve against natural selection that he invoked group selection as a potential explanatory mechanism (Van Valen 1971). It is now well accepted that kin selection – the avoidance of kin-competition – can drive dispersal evolution (e.g., Gandon 1999), but it is also clear that dispersal evolves even in unrelated groups. Early models of dispersal evolution showed that, where demes differed in carrying capacity, dispersal should evolve to be zero because the majority of dispersing individuals end up moving to less favourable habitat (Balkau and Feldman 1973; Hastings 1983; Holt 1985). This ubiquitous cost of dispersing in a spatially heterogeneous landscape must be mitigated for dispersal to evolve beyond zero. While avoidance of kin competition is one mitigating benefit, it was also shown that dispersal is beneficial when temporal variation in habitat quality is present (e.g., Levin et al. 1984; Moody 1988; McPeek and Holt 1992). Thus, while spatial heterogeneity alone would drive dispersal rates down, temporal heterogeneity drives dispersal rates upwards.

A nice exemplar of spatiotemporal variation in habitat lies the metapopulation, in which demes experience stochastic extinction. Because empty patches can only be colonised by dispersers, not dispersing will eventually result in extinction, and so we expect non-zero dispersal rates to evolve despite costs to individual fitness. The observation that evolutionary pressures on dispersal might maintain fitness costs against natural selection was sufficiently noteworthy that the effect was named “the metapopulation effect” (Olivieri et al. 1995).

### The discovery of spatial sorting

Since work on the metapopulation effect, an even more striking case of dispersal evolution has become apparent: the evolution of dispersal on invasion fronts. In this situation, a population is spreading into uncolonised habitat, and the individuals on the very edge of this invasion can only be there because they have dispersed further than their conspecifics. This sets up conditions for assortative mating by dispersal ability each generation, leading to the runaway evolution of dispersal on an invasion front. Thus, on invasion fronts, we see directional evolution of traits related to dispersal, but the mechanism by which this occurs is not natural selection; directional evolution occurs despite the absence of a fitness differential. These observations suggest that natural selection may have a “shy younger sibling” – dubbed “spatial sorting” – and that spatial sorting may be a spatial analogue of natural selection (Shine et al. 2011). Spatial sorting arises when there is a cline in density and variation for dispersal ability. If true, this would make spatial sorting an important mechanism by which dispersal evolves in temporally heterogeneous populations and, potentially, the primary cause of the metapopulation effect. The idea of spatial sorting was first adumbrated by Cwynar and MacDonald (1987) and has been re-discovered and refined by various authors since (e.g. Travis and Dytham 2002; Hughes et al. 2003; Phillips et al. 2008; Burton et al. 2010; Shine et al. 2011).

In many contexts, spatial sorting is likely so fleetingly transient as to be easily missed. In the context of spatially expanding populations, however, spatial sorting is persistent on the invasion front for the duration of spread and for some time after. In this situation, the invasion front colonizes unoccupied space every generation, spatial sorting is sustained over time on the invasion front, and its effects are difficult to ignore. The clearest natural example of this comes from the spread of cane toads across northern Australia. Here, evolved shifts in dispersal ability contributed to a five-fold increase in invasion speed over 70 generations (Phillips et al. 2008, 2010; Perkins et al. 2013). Numerous other natural examples have come to light, ranging across taxa from insects to plants (e.g., Cwynar and MacDonald 1987; Simmons and Thomas 2004; Lombaert et al. 2014). More compelling still, are a growing list of laboratory studies showing repeatable evolutionary shifts in dispersal on invasion fronts (e.g., Ditmarsch et al. 2013; Fronhofer and Altermatt 2015). Two recent laboratory studies on beetles also unequivocally demonstrate that these evolutionary shifts are due to spatial sorting (Ochocki and Miller 2017; Weiss-Lehman et al. 2017).

### The theory of spatial sorting

The original theoretical arguments behind spatial sorting on invasion fronts were illustrated with individual-based simulation models (e.g., Travis and Dytham 2002; Hughes et al. 2003; Phillips et al. 2008; Burton et al. 2010; Shine et al. 2011). While these painted a convincing picture and pointed to an interesting range of theoretical possibilities (including the possibility that spatial sorting could act aginst natural selection, causing a reduction in fitness), those findings are difficult to generalise. More recently, theoreticians have worked to integrate variation in dispersal into equation-based models of biological spread. Although mathematically challenging, these equation-based models have the potential to generate cleaner notions of how certain biological factors modulate the dynamics of spatial sorting. These models treat dispersal as a quantitative trait embedded within an integral projection (e.g., Ellner and Schreiber 2012), integrodifference (e.g., Perkins et al. 2013), or partial differential (e.g., Perkins et al. 2016) model. Additionally, an important generalisation of Fisher’s (Fisher 1937) reaction-diffusion model has been conceived that elegantly encapsulates the process of spatial sorting by allowing the model’s “diffusion coefficient”, *D*, to evolve (Benichou et al. 2012; Bouin et al. 2012; Bouin and Calvez 2014). This new class of reaction-diffusion model is currently the focus of intense and productive theoretical work, much of which necessarily involves advanced mathematics and is beyond the grasp of many biologists.

Because spatial sorting was first discovered in the context of invading populations, the mathematical treatment of this process has utilised the tools of that field, primarily reaction diffusion equations and integro-difference models. Likewise, much focus has been placed on the emergent property that dispersal evolution leads to accelerating invasion fronts. In this context, the primary focus of has been on the evolution of equilibrium states of dispersal traits. Substantially less attention has been paid to the mechanism itself, spatial sorting: how it might be described, and the conditions under which it operates. This is an omission, because precise articulation of the mechanism sharpens our intuition about the process and enables deeper theoretical work, such as that which has been built on formal mathematical descriptions of natural selection. Here, we focus exclusively on the mechanism of spatial sorting, and we explore how it might be expressed in terms of classical population genetic models.

### A theoretical reorientation

We start with the basic haploid and diploid models of natural selection (Crow and Kimura 1970). These models, originally conceived free of population dynamics and expressed as either a change in allele ratios or allele frequencies over time (Haldane 1924; Wright 1931), have become the standard models by which students are introduced to the theory of natural selection (e.g., Hartl et al. 1997) and over the years have become more explicitly linked to population dynamics (e.g., Crow and Kimura 1970). A particularly thorough and lucid derivation is given in Otto and Day (2007), in which we see the frequency of the *A* allele, *p*, following births and deaths given by

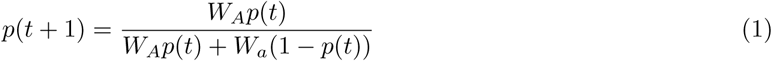

for a haploid system, and

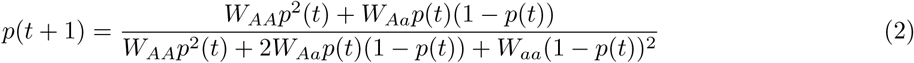

for a diploid system. In these models, the *W* terms represent the fitnesses (per capita balance of births and deaths) of individuals carrying each genotype, and the denominator in each case is the mean fitness of the population.

In this paper, we introduce a simple conceptual arrangement that allows us to introduce spatial sorting into these foundational models of evolutionary biology. We demonstrate that, on an invasion front, spatial sorting can be conceptualized precisely as selection operating through space, rather than time, and that this conceptualization leads to a simple generalisation of the haploid and diploid selection models. This generalisation recognises that fitness can have both temporal and spatial aspects, and it is this spatiotemporal fitness that is maximised on an invasion front. The maximisation of spatiotemporal fitness, rather than classical temporal fitness, explains why dispersal rate evolves upwards on invasion fronts, but also explains why this can come at a cost to classical fitness. Our simple models are easy to understand and, like the classic models of natural selection, will provide a launching pad for many theoretical forays in population genetics.

## The conceptual arrangement

Space and time are discrete. We imagine a one-dimensional spatial lattice of large size, with position on the lattice denoted by *x* (see Fig. 1). At a given time *t*, birth, death, and dispersal occur, in this order.

**Figure 1:**
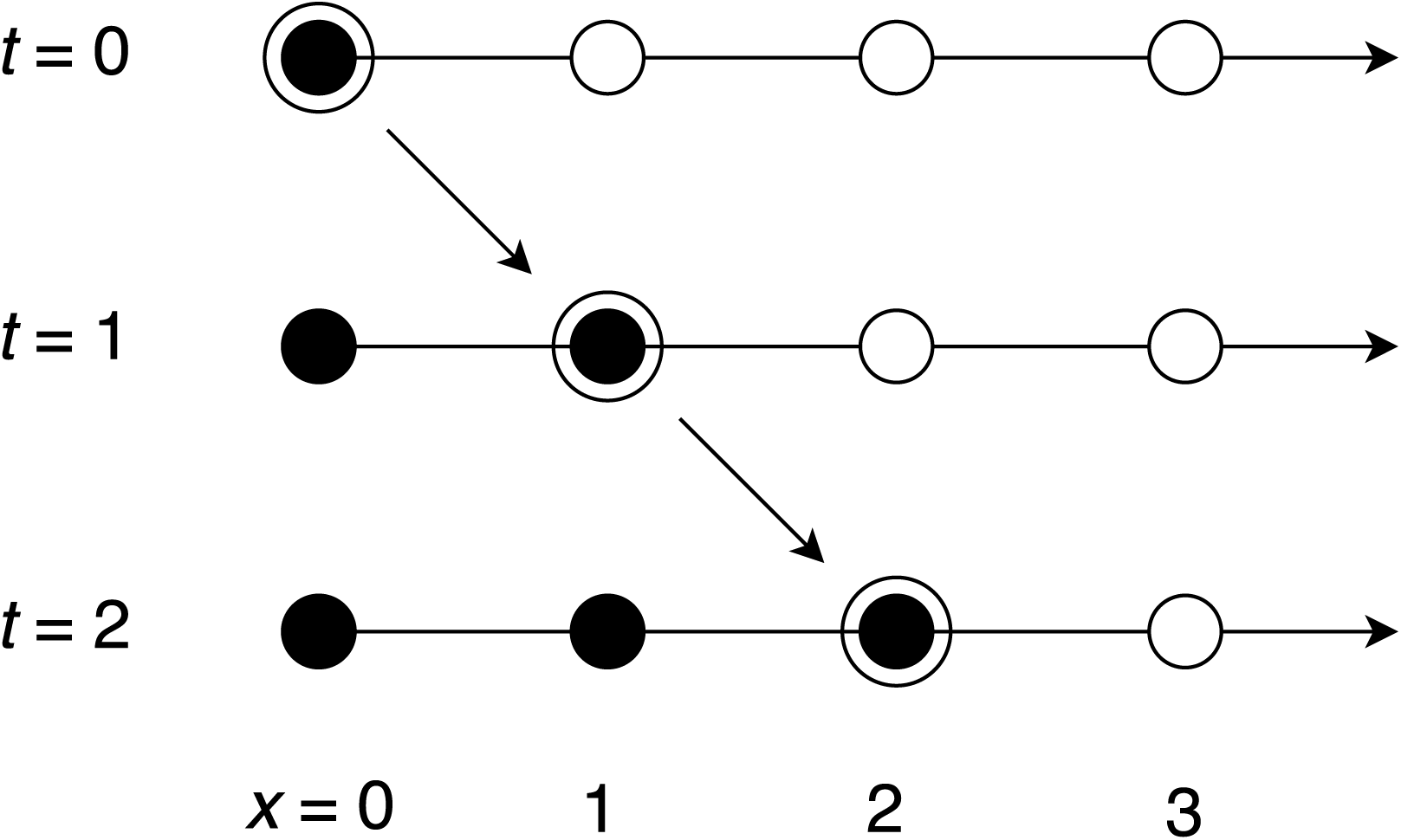
The spatiotemporal arrangment of the model. Filled black circles show space being occupied as time progresses. The model tracks the dynamics of the foremost patch at *x* = *t*, shown by the large outer circles and the diagonal arrows.

At *t* = 0, we imagine that only the left-most patch in our lattice, at *x* = 0, is occupied, containing a large number of individuals, *n*. All other patches are empty. We assume nearest-neighbor dispersal, which implies that if the dispersal rate is not zero, patch 1 will be occupied at time 1, patch 2 at time 2, and so on. The model tracks the dynamics of this vanguard population as it moves through space, at *x* = *t* (Fig. 1). As long as the product of the mean population growth rate and mean dispersal rate is greater than 1, then this frontal deme will increase in size over time.

### Haploid dynamics

In a haploid model with discrete generations, we imagine two alleles, *A* and *a*. Each allele has both a spatial and temporal aspect to its fitness. *W*_*i*_ denotes temporal fitness of allele *i ∈ {A, a}*: the per capita number of surviving offspring at *t* + 1, such that *W*_*i*_ = (1 + *b*_*i*_)(1 *-d*_*i*_), where *b*_*i*_ is the expected net replacement per capita (through births and survival of parents) in each time interval and *d*_*i*_ is the probability of death in each time interval. *V*_*i*_ denotes spatial fitness of allele *i*, which we define as the probability of dispersal to *x* + 1. The fraction 1 − *V*_*i*_ that do not disperse might either stay at location *x* or move to location *x -* 1, but these outcomes do not affect the dynamics at the range front.

Following reproduction but before dispersal, the number of individuals of each genotype at time *t* and vanguard patch *x* is

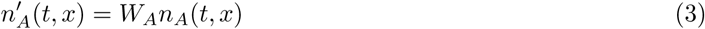

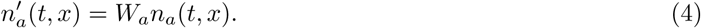

Following dispersal, the number of individuals of each genotype at time *t* + 1 and newly occupied patch *x* + 1 is

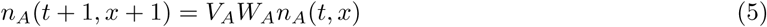

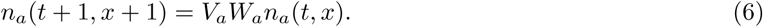

If we let 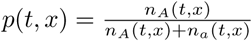, then

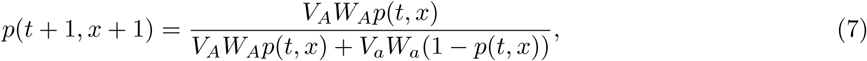

which is similar to the standard haploid model of selection (eqn. 1), except that we have explicitly incorporated *spatiotemporal fitness* via *V*_*A*_*W*_*A*_ and *V*_*a*_*W*_*a*_. In the event that temporal aspects of fitness are identical across the two alleles (that is, the two alleles have equal reproductive fitness), spatial fitness drives any evolutionary changes that occur; in the event that spatial fitness is identical between the two alleles (that is, the two alleles have equal dispersal tendency), we recover the standard haploid model of natural selection.

Similar to the standard model, eqn. 7 has two equilibria: at *p* = 0 (stable when 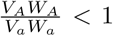) and *p* = 1 (stable when 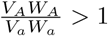). These stability criteria point to the strong interaction between temporal and spatial aspects of fitness, because we can think of them as products of two relative fitnesses, one spatial and one temporal. The implication of this result is that the vanguard will eventually be dominated by one genotype or the other, provided that spatial expansion continues for long enough.

Another clear implication of the stability criteria, however, is that spatial sorting can drive populations on the invasion front to lower fitness. Allele *A* will move to fixation as long as 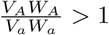. If the spatial fitness of *A* is sufficient to offset a temporal fitness cost, the invasion front evolves towards lower temporal fitness (see Fig. 2). It is clear that, on an invasion front, it is not temporal fitness that is being maximised, but spatiotemporal fitness.

**Figure 2:**
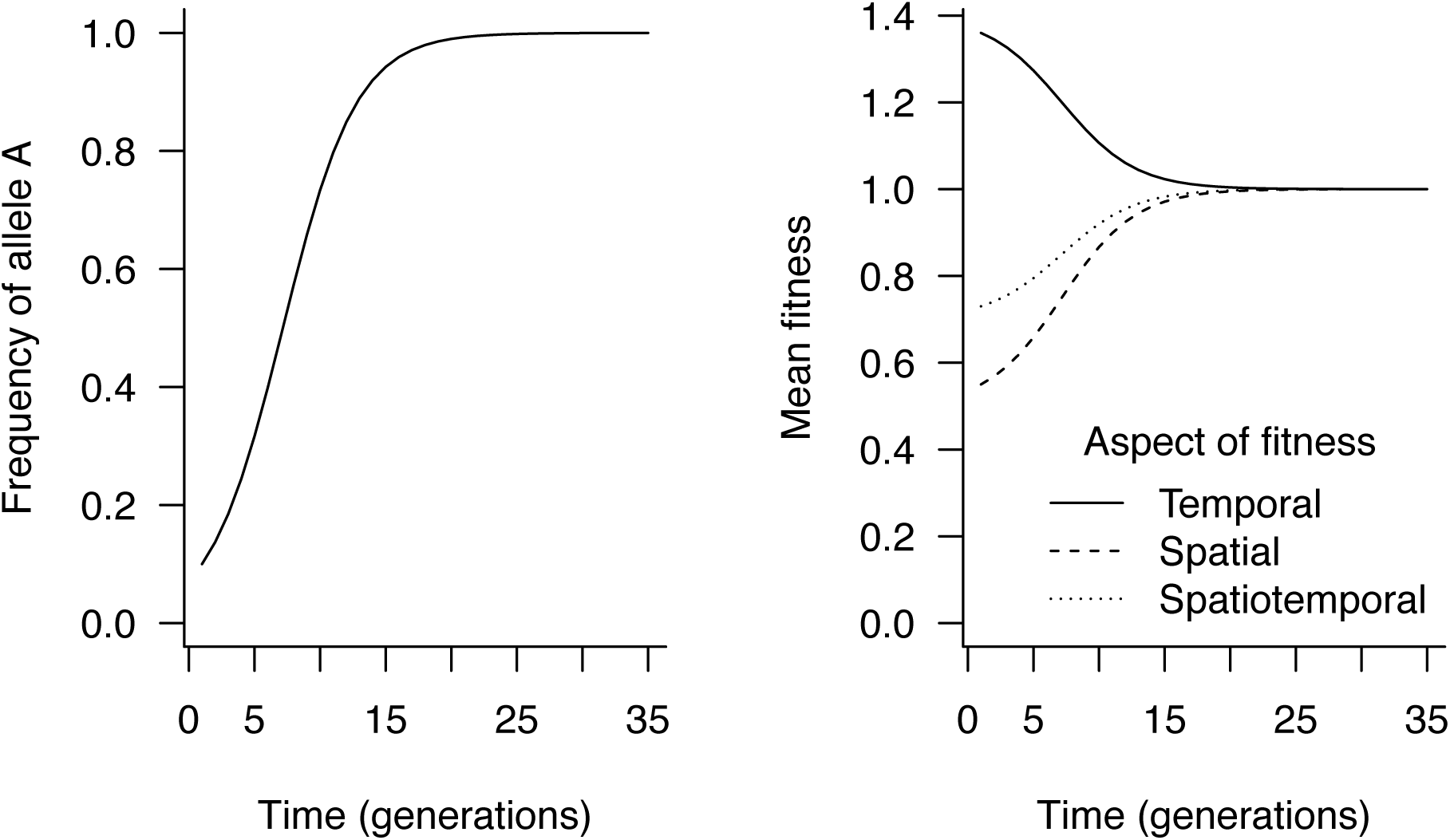
Haploid evolutionary dynamics on an invasion front. Here, the frequency of allele *A* increases because it has the higher spatial fitness, and this occurs despite it having substantially lower (temporal) fitness. The invasion front evolves to maximise spatiotemporal fitness rather than traditional fitness as measured by reproductive rate. Parameters: *W*_*A*_ = 1, *W*_*a*_ = 1.4, *V*_*A*_ = 1, *V*_*a*_ = 0.5.

### Diploid dynamics

In a diploid model we have three genotypes, *AA, Aa*, and *aa*, and a total population size *n*(*t, x*) = *n*_*AA*_(*t, x*) + *n*_*Aa*_(*t, x*) + *n*_*aa*_(*t, x*). Following birth, death, and dispersal, the numbers of the three genotypes at time *t* + 1 and patch *x* + 1 follow

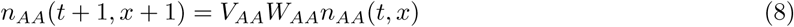

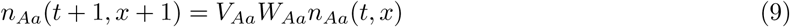

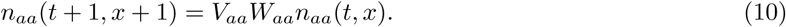

Again, we focus on the proportion, *p*, of *A* alleles in the population, which is defined as

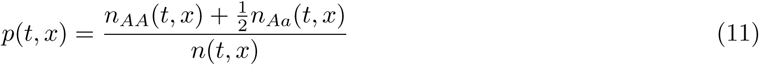

and at time *t* + 1 and patch *x* + 1 follows

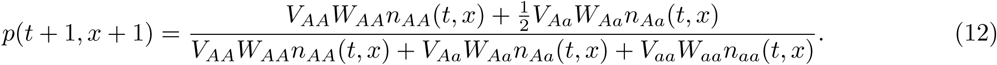

If we assume Hardy-Weinberg equilibrium and define *q* = 1 − *p*, then *n*_*AA*_ = *np*^2^, *n*_*Aa*_ = 2*npq*, and *n*_*aa*_ = *nq*^2^. Many well-known departures from this assumption are possible, with the magnitude of these departures influenced by the magnitude of differences among genotypes with respect to *V* and *W.* Nonetheless, these departures are, in practice, often small and have a tendency to vanish as time progresses. Assuming that departures from Hardy-Weinberg equilibrium are modest, we can approximate the dynamics of *p* recursively according to

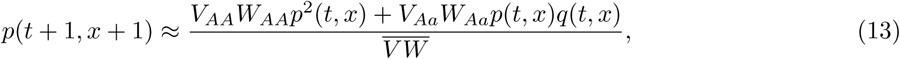

where 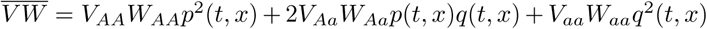 is the *mean spatiotemporal fitness* of the population. Again, this model is similar to the standard diploid model of selection (eqn. 2), except that fitness is now explicitly spatiotemporal. Also as before, in the event that there is no spatial fitness differential (*V*_*AA*_ = *V*_*Aa*_ = *V*_*aa*_), the model reduces to the standard diploid model (eqn. 2).

This model has three equilibria. The equilibria at *p* = 0 and *p* = 1 are stable when 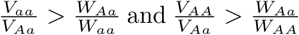, respectively. An equilibrium allowing for coexistence of the two alleles occurs at 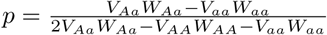 and is stable when both *V*_*Aa*_*W*_*Aa*_ > *V*_*AA*_*W*_*AA*_ and *V*_*Aa*_*W*_*Aa*_ > *V*_*aa*_*W*_*aa*_. It is noteworthy that, unlike the haploid case, the vanguard can be populated by a stable mixture of *A* and *a* alleles but that this requires some manner of trade-off between the life-history and dispersal traits of *AA* and *aa* genotypes that result in the *Aa* genotype having the highest spatiotemporal fitness.

As with the haploid model, conditions on the invasion front maximise spatiotemporal fitness rather than traditional fitness as measured solely by the balance of births and deaths. Figure 3 gives an example of these dynamics, showing a situation in which the *A* allele shows a dominant pattern of expression, and has higher spatial, but lower temporal fitness than the *a* allele.

**Figure 3:**
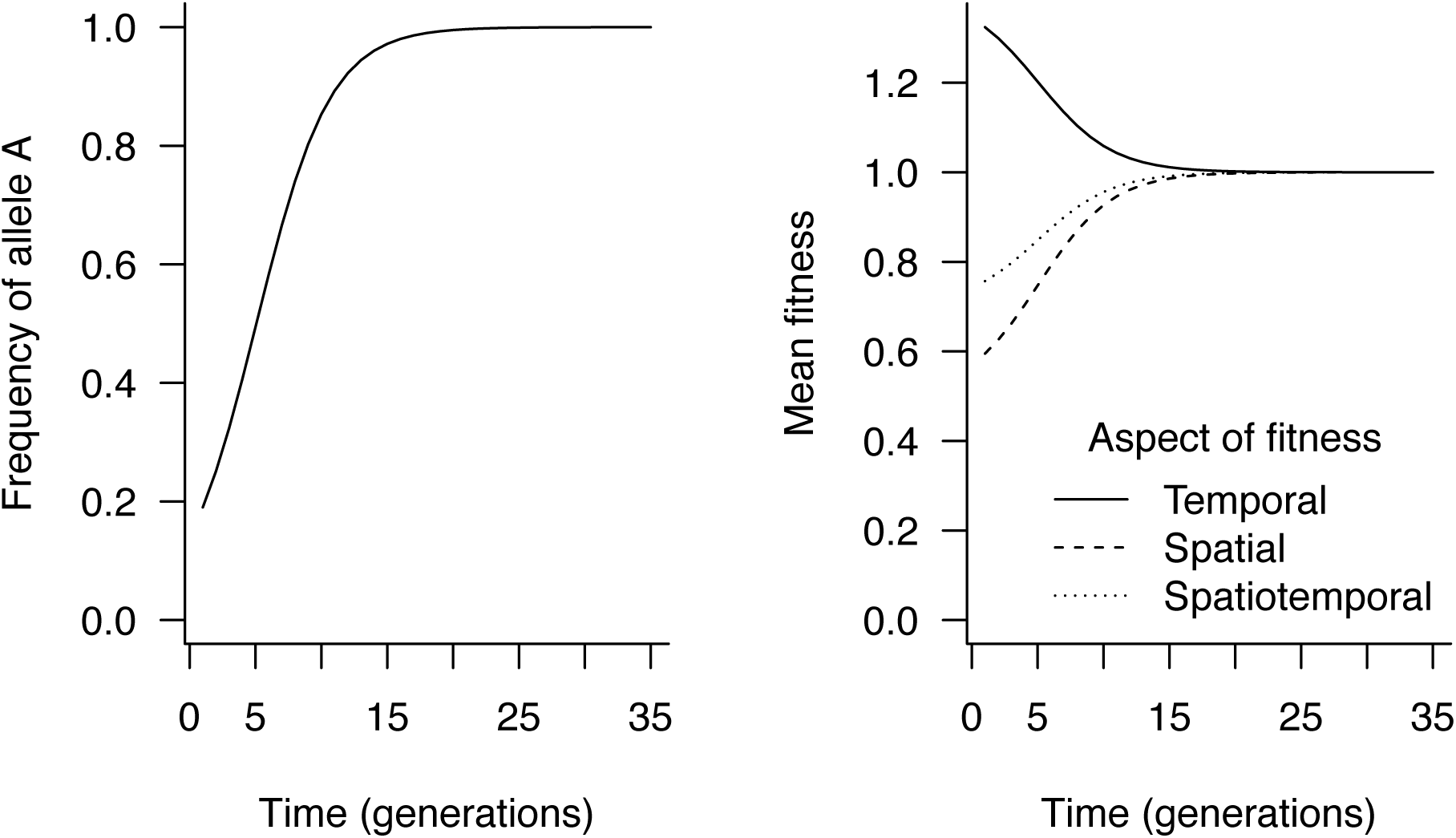
Diploid evolutionary dynamics on an invasion front, as described by equations (8)-(11). Here, the *A* allele shows a dominant expression pattern. The frequency of *A* increases because it has the higher spatial fitness, and this occurs despite it having lower (temporal) fitness than allele *a*. The invasion front evolves to maximise spatiotemporal fitness rather than traditional fitness as measured by reproductive rate. Parameters: *W*_*AA*_ = 1, *W*_*Aa*_ = 1, *W*_*aa*_ = 1.4, *V*_*AA*_ = 1, *V*_*Aa*_ = 1, *V*_*aa*_ = 0.5.

R Code implementing numerical examples of these recursions is available at https://github.com/benflips/spatSortNumerical.

## The effects of gene flow on the response to selection

It is well established that gene flow can undermine selection by contributing maladapted genes into a population. Spatial fitness only makes sense in models in which gene flow is a fundamental force. Because of this, it is worth considering how gene flow influences the response to spatial sorting, and comparing this to the classic result in which gene flow influences the response to natural selection. One way of conceptualising this is to use our moving frame of reference (as per the above model), but now imagine that the next patch (at *x* = 1) is already occupied by a population with given density, *n*_1_, and allele frequency, *p*_1_. In this frame of reference, we track the gene frequencies of our immigrants and consider “gene flow” to be effected by the resident population into which our immigrants arrive.

### Haploid occupied-patch model

In this situation, following reproduction and dispersal, the composition of individuals leaving patch 0 is identical to that given in equation 7, and the total number of these individuals is 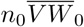. For simplicity, we ignore evolutionary and dispersal dynamics in patch 1 and simply allow it to be occupied by *n*_1_(*t*) individuals, with the frequency of *A* alleles in this patch given by *p*_1_(*t*). In this scenario,

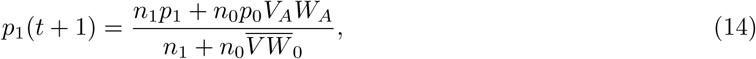

and the change in allele frequency in our moving window between (*t* = 0, *x* = 0) and (*t* = 1, *x* = 1) is

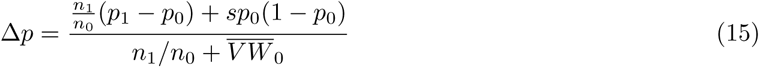

where *s* = *V*_*A*_*W*_*A*_ − *V*_*a*_*W*_*a*_ is the spatiotemporal selection differential.

This equation is similar to that generated under consider classic selection-migration balance (Wright 1931, 1940): certainly, the equilibrium conditions are identical to the classic case (with our 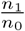 taking the place of Wright’s *m* in the classic model). Here, however, we have a moving frame of reference; we consider immigrants as our reference population, with the disruptive effect of “gene flow” provided by the resident population at *x* = 1. This is an unusual perspective to take, but is consistent with the previous model. It also allows us to group spatial and temporal fitness effects as the sum of changes due to gene flow (first term in the numerator of eqn. (15)) versus natural selection and spatial sorting (second term in the numerator of eqn. (15)).

As with classic selection-migration balance, the effect of gene flow in this scenario depends on the relative number of residents versus migrants, and the difference in allele frequencies between our two populations (Crow and Kimura 1970).

### Diploid occupied-patch model

Similar reasoning can be applied to derive a map for a diploid occupied-patch model, but here we have

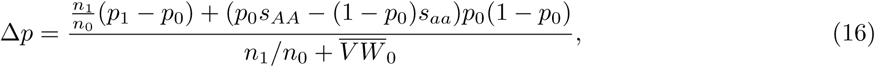

where *s*_*AA*_ and *s*_*aa*_ represent the difference in spatiotemporal fitness between each of the homozygotes and the heterozygote: *V*_*AA*_*W*_*AA*_ − *V*_*Aa*_*W*_*Aa*_ and *V*_*Aa*_*W*_*Aa*_ − *V*_*aa*_*W*_*aa*_, respectively.

## Discussion

By focusing only on the evolutionary dynamics of an invasion front, the invasion front model developed here gives natural selection’s shy younger sibling, spatial sorting, nowhere to hide (Shine et al. 2011). In the special context of the invasion front, spatial sorting is no longer a transient phenomenon—it is persistent through time as long as the front is expanding—and this allows us to see that spatial sorting is akin to natural selection. Whereas natural selection operates to filter genotypes through time, spatial sorting operates to filter genotypes through space. Whereas natural selection filters genotypes on the basis of reproductive rate, spatial sorting filters genotypes on the basis of dispersal rate. As a result, we are justified in thinking of genotypes having both temporal and spatial aspects to their fitness.

In the special case of an invasion front, spatial sorting happens every generation and so exerts influence similar to that exerted by traditional natural selection. On invasion fronts, it is clear that natural selection and spatial sorting interact strongly; a situation most clearly seen in the stability criteria of the models here. The stability criteria in all cases express an imbalance between relative spatial and temporal aspects of fitness. If the relative spatial fitness of allele *A* is greater than the relative temporal fitness of allele *a*, then *A* will increase in frequency, even if this entails a reduction in traditional (temporal) fitness. Conditions on the invasion front clearly maximise spatiotemporal fitness rather than traditional fitness based only on reproduction and survival. On the invasion front we can see, with unusual clarity, why dispersal might evolve at the expense of fitness, and the mechanism is a spatial analogue of natural selection.

Because dispersal is a necessary condition for gene flow to occur, most models of dispersal evolution must account for gene flow. By taking a moving frame of reference, our invasion front model experiences none of the homogenizing influence of gene flow. On the invasion front, we can simplify the spatial problem by making it unidirectional, like time. This approach allows us to observe spatial sorting at play because it eradicates gene flow. In this frame of reference, gene flow is provided by the population resident inside a patch prior to the arrival of our focal population. This perspective preserves the process of spatial sorting, but shows that spatial sorting is undermined by gene flow in precisely the same manner as natural selection.

Models that examine the evolution of dispersal typically focus on the evolutionary optima that emerge under various spatiotemporal scenarios. It is regularly observed that dispersal evolves, despite evident costs to mean fitness (Johnson and Gaines 1990). While kin-competition is one mechanism that explains this (that is, the loss of individual fitness is mitigated by a gain in inclusive fitness, e.g., Hamilton and May 1977), there is clearly a spatial mechanism at play also (e.g., Olivieri et al. 1995). Spatial sorting is almost certainly that mechanism, and so it is important that we develop models that describe and isolate it. To this end, invasion fronts make a powerful study system. But perhaps because spatial sorting has come to prominence in the context of biological invasions, the development of mathematical theory of spatial sorting has mostly bypassed simple mechanism-focussed models such as ours in favor of complex models that predict outcomes across all space (e.g., Benichou et al. 2012; Perkins et al. 2016). The intent of the formulation developed here is to go back to basics, focus carefully on the mechanism, and show how spatial sorting relates to basic evolutionary theory. The simple theoretical tools that we make use of are not new, of course. Population genetic models accounting for interactions between natural selection and gene flow, for example, go back decades (e.g., Nagylaki 1992). Such models have, however, yielded great theoretical insight and are also useful for applied topics, such as adaptation in response to a shifting environment (e.g., Case and Taper 2000). Thus, linking spatial sorting to basic population genetics connects it to a literature that is already rich in biological detail and steeped in demonstration of applied value.

While our model very simply captures the process of spatial sorting, it is limited. One clear limitation is that we are not focussed on invasion fronts as a phenomenon, necessarily. Models of invasion fronts are often concerned with estimating spread rate, for example, and this requires tracking the rate at which a threshold density moves through space. Our frame of reference is not defined by a particular population density, so our model cannot track spread velocity in this way. Despite this limitation, our finding that the product of spatial and temporal fitnesses is maximised has intriguing links to spread theory. Classic spread theory (in continuous space and time) shows that spread rate, 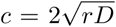, where *r* is the rate of increase of the population, and *D* is the diffusion coefficient, defining dispersal rate (Fisher 1937; Skellam 1951). To the extent that our *W* maps to *r* and our *V* maps to *D*, our result suggests that conditions on the invasion front select for whatever increases invasion speed. A similar argument has recently been made by by Deforet *et al.* (2017) using a reaction diffusion framework with competing clonal lines. Thus, while the link between selection, sorting, and spread rate is an aspect requiring further theoretical development, there is a strong hint from both of these results that evolution maximises the spread rate of an expanding population by maximising spatiotemporal fitness on the invasion front.

Importantly, this maximisation of spatiotemporal fitness will occur even if this comes at a cost to traditional fitness (the balance of births and deaths). Thus, spatial sorting is a directed evolutionary process that, in particular circumstances, can act to reduce the traditional, temporal fitness of a population. On invasion fronts, it is now well established theoretically that fitness will often be reduced by the serial foundering that occurs as a population spreads (Slatkin and Excoffier 2012; Peischl et al. 2013) and this “expansion load” can slow invasions (Peischl et al. 2015; Phillips 2015). Our results show a deterministic process that can also erode fitness – spatial sorting – but unlike the stochastic case, our deterministic process can only erode fitness if doing so increases spatiotemporal fitness. Thus, the deterministic process we describe can only cause invasions to accelerate. The tight interaction between spatial and temporal fitness in our model also has implications for how covariation between dispersal and fitness will constrain trait evolution on invasion fronts. For example, in our model, if a mutant allele confers an increase in fitness but also causes a proportional reduction in dispersal, spatiotemporal fitness remains unchanged and we would expect this mutant to have identical spatiotemporal fitness to the wild-type. Thus, the scaling between spatial and temporal aspects of fitness has important implications for how evolution might proceed on invasion fronts; an area worthy of further exploration. Under more complicated models, interactions between the temporal and spatial components of fitness can become even richer, as revealed in a model of cane toad spread that pointed to increased temporal fitness giving rise to a larger spatiotemporal fitness differential under density dependence (Perkins et al. 2013).

More generally, assuming that a gene has the potential to jointly affect both spatial and temporal aspects of fitness may well prove useful. In many real cases, the same trait can affect both dispersal and survival/reproduction (Burgess et al. 2016). These dual purposes cause substantial ambiguity as to why a trait may have evolved: was it for dispersal, or fitness? If we accept the view that spatiotemporal fitness is a useful concept (if for no other reason than obviating this duality), then the moving frame of reference approach we use here might profitably be applied elsewhere. Classical population genetics very sensibly focusses on a population at a particular location. In this setting, immigration (from another location) acts to disrupt the process of adaptation and push populations away from their local optimum, contributing a “migrant load” of non-locally-adapted alleles to the population. In a moving frame of reference, we instead focus on the migrant population, and treat the resident population as the agent of gene flow. The classic view is often simplified by assuming that migration (and selection) are small forces; a view that is well justified in populations close to some demographic and evolutionary equilibrium (Crow and Kimura 1970). In cases where migrants are actually the bulk of a population, however, the moving frame of reference approach makes more sense: we can again assume that gene flow (effected by residents in this case) is a small force. The fact that we can (with the moving frame of reference) derive essentially the same equations for migration-selection balance as the classic case (Wright 1931) suggests that there may be much theoretical symmetry between these two alternate views of the population.

On the whole, our results speak to the striking similarity of action between natural selection and spatial sorting on an invasion front. The analogy we strike is similar to that of Slatkin and Excoffier (2012), who show that serial founder events on an invasion front are the spatial analogue of genetic drift. In both cases, we see space playing the role of time in classic evolutionary models. This similarity hints that many theoretical results—built off the analogous standard models that underlie much of evolutionary theory—may also apply on invasion fronts subject to spatial sorting alone. That is, if there is no difference between alleles in aspects of temporal fitness (*W* terms are all equal), then the equations become identical to the standard haploid and diploid models, but they refer to spatial sorting rather than natural selection. Thus, we might expect that many theoretical results extending the basic model—exploring issues of dominance, frequency dependence, sexual conflict, drift vs selection, and so on—can be re-derived for spatial sorting. Indeed, by combining Slatkin and Excoffier’s (Slatkin and Excoffier 2012) approach to serial foundering with our notion of spatial sorting, it has already proved possible to capture the drift/selection dynamics of dispersal-modifying alleles on invasion fronts (Peischl and Gilbert 2018). Through this or other avenues, our simple formulation has the potential to provide a useful entry point into fertile theoretical ground for some years to come.

## Acknowledgements

The manuscript was improved by thoughtful comments from Stephan Peischl, Susan Frances Bailey, and Frédéric Guillaume. BLP was supported by an Australian Research Council Future Fellowship (FT160100198).

